# Computational metabolism modeling predicts risk of distant relapse-free survival in breast cancer patients

**DOI:** 10.1101/468595

**Authors:** Lucía Trilla-Fuertes, Angelo Gámez-Pozo, Mariana Díaz-Almirón, Guillermo Prado-Vázquez, Andrea Zapater-Moros, Rocío López-Vacas, Paolo Nanni, Pilar Zamora, Enrique Espinosa, Juan Ángel Fresno Vara

**Author notes:** These authors contribute equally to this work (Lucía Trilla-Fuertes, Angelo Gámez-Pozo). Corresponding author e-mails: LT-F; AG-P; MD-A; GP-V; AZ-M; RLV; PN; PZ; EE; JAFV.

## Abstract

**Aims:** Differences in metabolism among breast cancer subtypes suggest that metabolism plays an important role in this disease. Flux Balance Analysis is used to explore these differences as well as drug response.

**Materials & Methods:** Proteomics data from breast tumors were obtained by mass-spectrometry. Flux Balance Analysis was performed to study metabolic networks. Flux activities from metabolic pathways were calculated and used to build prognostic models.

**Results:** Flux activities of vitamin A, tetrahydrobiopterin and beta-alanine metabolism pathways split our population into low- and high-risk patients. Additionally, flux activities of glycolysis and glutamate metabolism split triple negative tumors into low- and high-risk groups.

**Conclusions:** Flux activities summarize Flux Balance Analysis data and can be associated with prognosis in cancer.

## Introduction

Breast cancer has a high incidence, with 266,120 estimated new cases and 40,920 estimated deaths in women in the United States during 2018 [1]. The expression of estrogen receptor, progesterone receptor and human epidermal growth factor receptor 2 (HER2) classifies breast cancer into one of three groups: hormone receptor positive / HER2 negative (ER+), HER2 positive (HER2+) or triple negative (TNBC). In previous works, we defined a new subtype within ER+ tumors with a clinical outcome and molecular features more similar to TNBC. This new subtype was called TN-like. ER+ tumors which still have ER+ characteristics were renamed as ER-true [2].

Reprogramming of metabolism is a hallmark of cancer [3]. Tumor cells use glucose to produce lactate, thus avoiding glucose metabolism through the Krebs cycle [4]. Tumor cells also produce lactate from glutamine and then generate NADPH in a process known as glutaminolysis [5]. However, not all tumors show the same metabolic alterations. In previous studies we described that there are differences in metabolism between breast cancer subtypes [6].

Proteomics provides detailed information about biological processes. Technical improvements currently allow the quantification of thousands of proteins. This information can be used to calculate the output of metabolic pathways with Flux Balance Analysis (FBA). FBA is a computational method used to study metabolic networks, making possible to predict tumor growth rate or the rate of production of a metabolite [7]. In our previous study, preliminary results of FBA were shown. FBA predicted a higher growth rate in TN-like tumors than in ER-true. Moreover, tumor growth predictions for TNBC and TN-like were comparable. We have proposed the calculation of flux activities as a summary measurement of flux distributions and showed that flux activities can be used to compare metabolic patterns between tumors or cells [8].

The aim of this work is to study in depth the metabolic differences previously characterized in breast cancer [2,6]. More specifically, proteomics data were analyzed through FBA to find metabolic pathways with prognostic value.

## Materials and Methods

### Patient cohort

One hundred and six formalin-fixed paraffin-embedded (FFPE) samples from I+12 Biobank and from IdiPAZ Biobank, both integrated in the Spanish Hospital Biobank Network (RetBioH; www.redbiobancos.es), were analyzed. This study was approved by the Ethical Committees of Hospital 12 de Octubre and Hospital La Paz.

Samples were selected according to these criteria: 1) node-positive disease, 2) no HER2 overexpression, and 3) patients treated with adjuvant chemotherapy and hormonal therapy in the case of ER+ tumors. The following clinical data were recorded: patient’s age, tumor size, lymph node status, tumor grade, adjuvant therapy administered and distant metastasis-free survival.

### Protein isolation

Proteins were isolated from FFPE samples as described in previous works [6,9]. Briefly, FFPE slices were deparaffinized in xylene and washed with absolute ethanol. Extracts were prepared in 2% SDS buffer using a protocol based on heat-induced antigen retrieval [10]. Protein concentration was determined using the MicroBCA Protein Assay Kit (Pierce-Thermo Scientific). Then, protein extracts were digested by trypsin and SDS was removed. Samples were dried and resolubilized in 15 µL of a 0.1% formic acid and 3% acetonitrile solution.

### Mass-spectrometry experiments and protein identification

Samples were analyzed on a LTQ-Orbitrap Velos hybrid mass spectrometer (Thermo Fischer Scientific, Bremen, Germany) coupled to NanoLC-Ultra system (Eksigent Technologies, Dublin, CA, USA). Peptides were separated on a self-made column. Solvent composition was 0.1% formic acid for channel A, and 0.1% formic acid and 99.9% acetonitrile for channel B. The mass-spectrometer was in data-dependent mode, acquiring full-scan spectra at a resolution of 30, 000 at 400 m/z after accumulation to a target value of 1,000,000, followed by CID (collision-induced dissociation) fragmentation on the twenty most intense signals per cycle.

The acquired data was analyzed by MaxQuant (version 1.2.7.4), and protein identification was done using Andromeda. Oxidation (M), deamidation (N, Q) and N-terminal protein acetylation was set as modifications. Protein abundance was calculated based on the normalized spectra intensity (LFQ).

Proteomics data were transformed into log2 and missing values were imputed to a normal distribution using Perseus [11]. Additionally, quality criteria as presence in at least 75% of the samples of each type or at least two unique peptides were required.

Proteomics data is publicly available in Chorus repository (http://chorusproject.org) under the name “Breast Cancer Proteomics”.

### Flux Balance Analysis and flux activities

FBA was performed using COBRA Toolbox available for MATLAB and the whole human metabolic reconstruction Recon2 [12,13]. The biomass reaction proposed in the Recon2 was used as an objective function to quantify tumor growth rate. Gene-Protein-Reaction rules (GPR), which relate genes and proteins with the enzymes involved in the reactions contained into the Recon2, were solved using a modification of the algorithm by Barker et al. [14]. Briefly, “OR” expressions are treated as a sum and “AND” expressions are calculated as the minimum [2,8]. Then, GPR values were introduced into the model using a modified E-flux consisted on normalize GPR data using a normal distribution formula [15].

Flux activities were calculated as in previous works [8]. Briefly, the flux activities were the sum of fluxes for all the reactions included in a metabolic pathway defined in the Recon2.

### Statistical analyses

Distant metastasis-free survival was selected as the prognostic variable. Statistical analyses were performed using GraphPad Prism v6 and predictors for distant metastasis-free survival was built using BRB Array Tool, developed by Dr. Richard Simon’s team. Cox regression models were done in SPSS IBM Statistics v20.

## Results

### Patient characteristics

One hundred and six samples were selected for the study. Finally, ninety-six patients diagnosed of breast cancer were analyzed. This cohort had been used in previous works from our group [2,6].

After performing the analyses of the 96 samples, of the 71 ER+ samples, 50 were reclassified as ER-true and 21 as TN-like, and twenty-six were TNBC tumors [2].

### Proteomics experiments

One hundred and six patients were recruited in this study. Four samples did not have enough protein amounts to perform mass-spectrometry (MS) experiments. After MS experiments, ninety-six samples had useful protein expression data. 3,239 proteins were measured. Quality criteria and filters were applied and a total of 1,095 proteins had two unique peptides and were detected in at least 75% of the samples of one type, i.e. ER+ or TNBC.

### Flux activities

FBA is a computational method used to study metabolic networks. The whole metabolic human reconstruction Recon2 was used to perform these analyses. The Recon2 is composed by 7,440 reactions and 5,063 metabolites grouped in 101 different metabolic pathways.

Using the results from FBA, flux activities were calculated as the sum of fluxes for the reactions included in each metabolic pathway described in the Recon2. Comparing breast cancer subtypes, significant differences in nucleotide interconversion pathway and reactive-oxygen species (ROS) detoxification were found between ER-true and TNBC and between ER-true and TN-like respectively (Figure 1).

**Figure 1:**
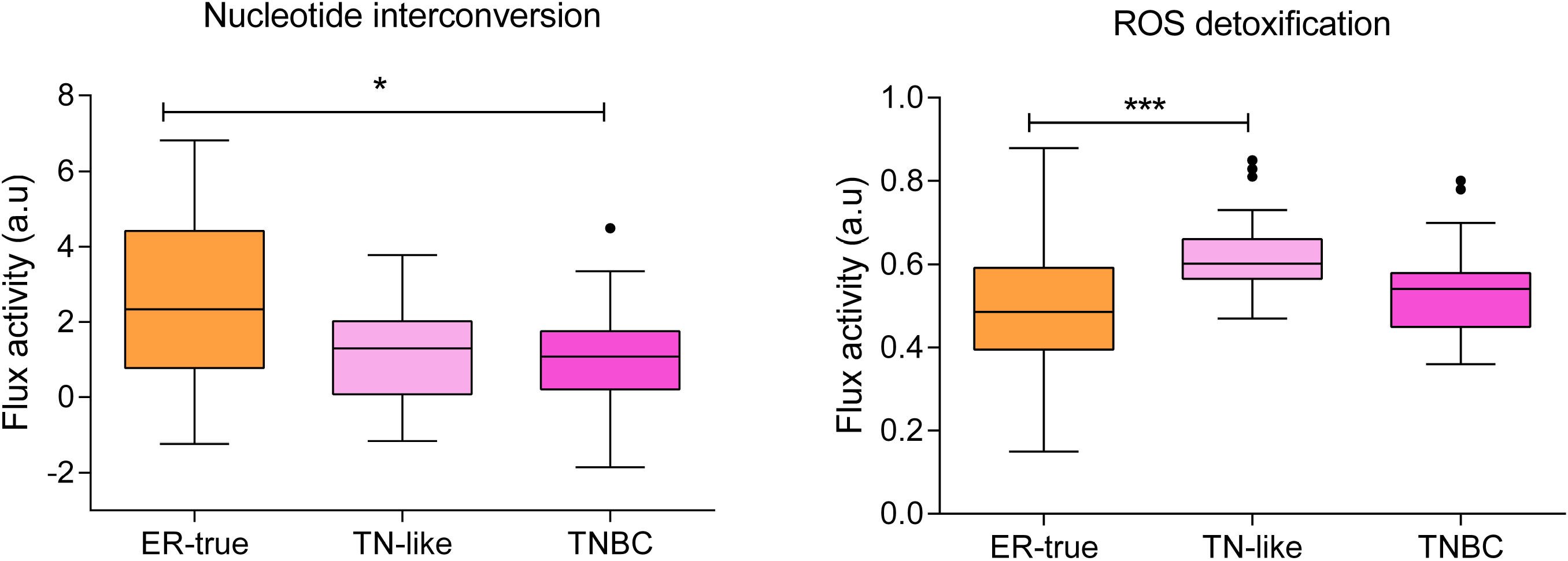
Flux activities with significant differences between breast cancer subtypes (* = 0.01 to 0.05, significant; *** = 0.0001 to 0.001, extremely significant).

### Prognostic signatures using flux activities

In order to identify flux activities related with distant relapse-free survival, BRB Array Tool from Dr. Richard Simon was used. We found five flux activities related with distant relapse-free survival (Sup Table 1).

Using these flux activities, a predictor of distant metastasis-free survival was constructed. This signature included the flux activity of three different pathways: vitamin A metabolism, tetrahydrobiopterin metabolism and beta-alanine metabolism. The predictor split our population into high-risk and low-risk groups (p-value = 0.0032, HR=6.520, cut-off= 30%-70%) (Figure 2, Table 2).

**Figure 2:**
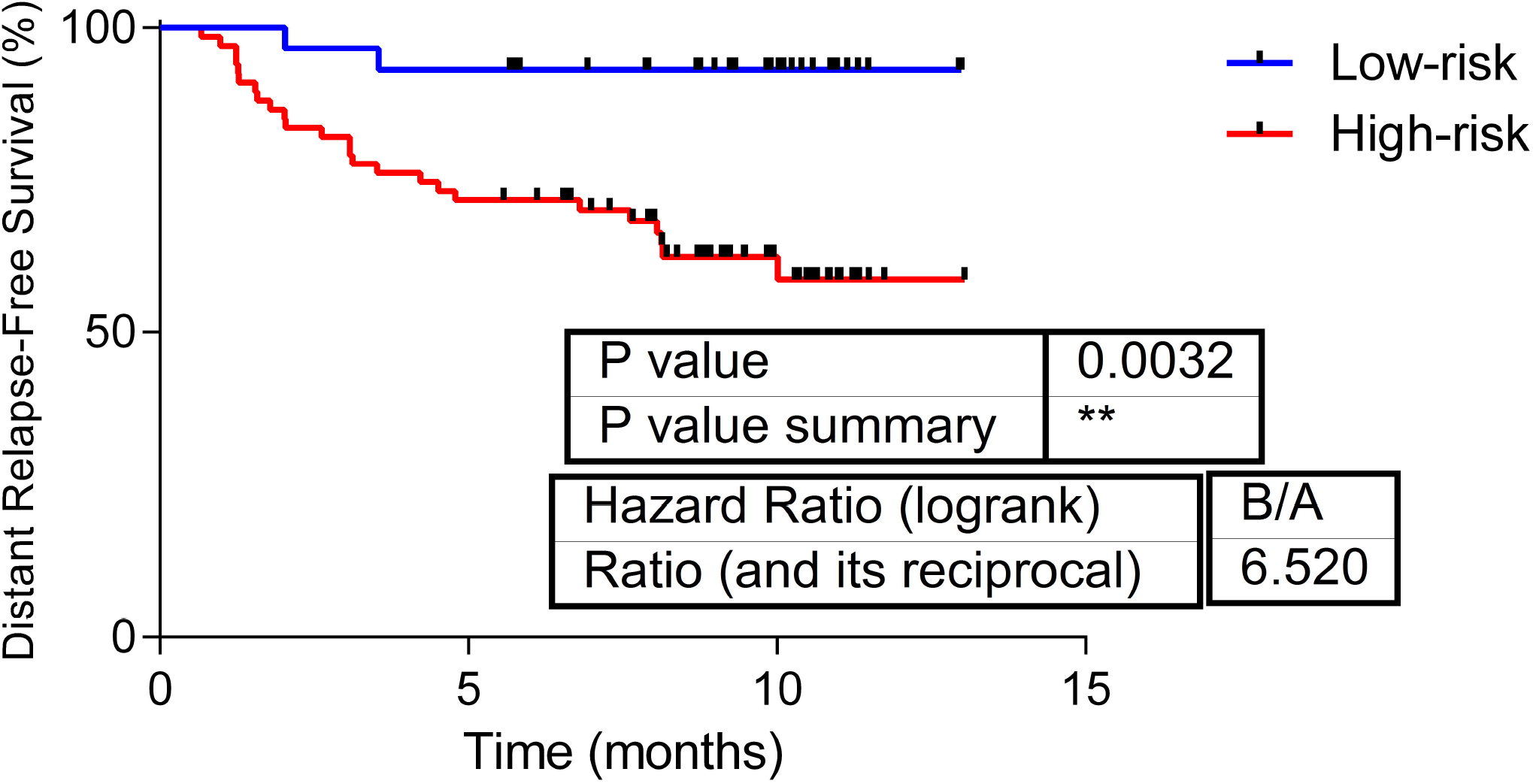
Predictive signature based on the flux activities of vitamin A metabolism, tetrahydrobiopterin metabolism and beta-alanine metabolism.

Strikingly, the prognostic signature retained its prognostic value in the ER+ tumors, being the differences statistically significant in the ER-true group (p-value= 0.0179), but have worse performance in TNBC tumors (Figure 3).

**Figure 3:**
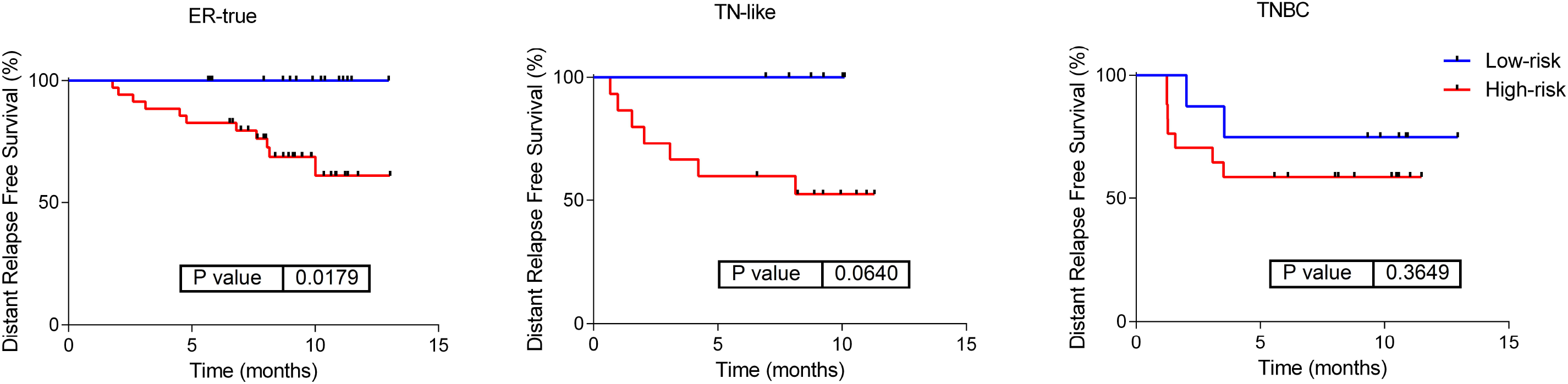
Predictor based on beta-alanine, tetrahydrobiopterin and vitamin A flux activities in the different breast cancer subtypes.

Multivariate Cox regression model showed that this signature added information to clinical data (Sup Table 2).

Given that TNBC tumors have differences in metabolism when compared with ER+; a predictive signature only for TNBC was built. This signature was formed by the flux activities of glycolysis and glutamate metabolism and split TNBC population into a low- and a high-risk group (p-value= 0.1064, HR= 4.600, cut-off= 30-70%); however, it did not reach significance (Figure 4, Table 3).

**Figure 4:**
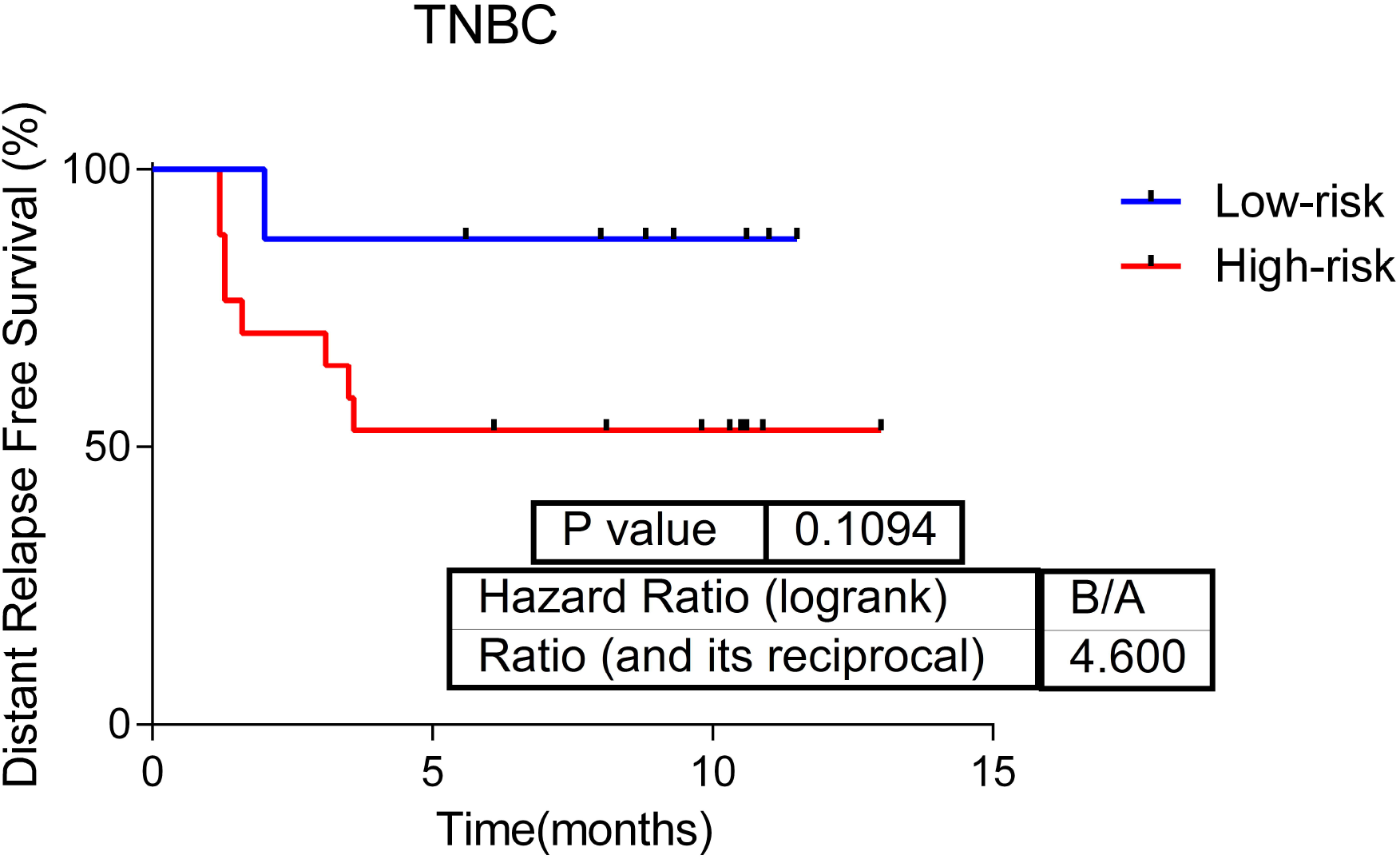
Predictor of distant relapse-free survival in TNBC tumors based on the flux activities of glycolysis and glutamate metabolism.

In this case, multivariate analysis was not significant (Sup Table 3).

## Discussion

Alterations in metabolism constitute a hallmark of cancer [3]. We previously described differences in metabolism between breast cancer subtypes [2,6]. TNBC and TN-like tumors had a higher growth rate than ER-true tumors in FBA predictions [2]. In the present work, our aim was to characterize additional differences in metabolic pathways between breast cancer subtypes.

The analysis of metabolism can use data coming both from genomics or proteomics. Proteomics provides more direct information about biological effectors than genomics, what is more useful in the GPR estimation. Proteomics experiments detected 3,239 proteins in the present study, although strict filtering criteria reduced the number to 1,095 proteins for analysis.

Comparing flux patterns in large cohorts is challenging. Flux activities are a method previously proposed to compare metabolic pathways between control cells and cells treated with drugs targeting metabolism [8]. Now, using flux activities we characterized differences in ROS detoxification and nucleotide interconversion pathways among breast tumor subtypes. Oxidative stress has been related with tumor aggressiveness in ER+ tumors [16]. TNBC tumors also have a high presence of ROS [17]. As expected, our model predicted lower ROS detoxification in ER-true tumors. Nucleotide interconversion category comprehends all the information about ATP and GTP metabolism. Differences in ATP or GTP production between breast cancer subtypes have not been previously described in the literature.

It was also possible to associate these flux activities with distant metastasis-free survival in this cohort. We found some flux activities related with distant relapse-free survival and built a predictor. The predictive signature included the flux activity of three metabolic pathways: vitamin A, tetrahydrobiopterin and beta-alanine metabolism. To our knowledge, this is the first time that data from FBA are associated with prognosis in cancer. Vitamin A or retinol has been related with risk of relapse in breast cancer [18], but there were no previous information about the prognostic value of tetrahydrobiopterin or beta-alanine metabolism. Additionally, the prognostic value of the predictor remained among ER+ tumors

It is well-known that TNBC tumors have differences in metabolism comparing with ER+ tumors [2,6]. For this reason, a predictor taking into account only TNBC tumors was done. This signature included flux activities of glycolysis and glutamate metabolism. However, survival and multivariate analyses are not significant, probably due to the small number of samples. An increase in glucose uptake and lactate production associated to glycolysis was previously described in TNBC cells [6] It is also well-known that lactate dehydrogenase B and other glycolytic genes are upregulated in TNBC tumors as compared with the other subtypes and that LDHB overexpression is associated with poor clinical outcome [19]. On the other hand, TNBC tumors have a deregulation of glutaminolysis and exhibited the most frequent expression of proteins related with glutamine metabolism than other subtypes [20,21].

The study has some limitations. A validation of the predictors in an independent cohort is needed. Another limitation is the number of samples of each group in this cohort. It is also necessary to validate these findings into larger cohorts of patients. On the other hand, further improvements in mass-spectrometry techniques now allow the detection of more proteins, which would provide further information at GPR level.

## Conclusions

In conclusion, flux activities, method proposed in previous works, now demonstrated its utility in summarizing FBA data and allows its association with prognosis. Differences in ROS detoxification and nucleotide interconversion pathways between breast cancer subtypes were characterized. Moreover, vitamin A, beta-alanine and tetrahydrobiopterin metabolism flux activities could be used to predict risk of distant metastasis-free survival in breast cancer patients. Finally, glycolysis and glutamate metabolism may be used to predict distant relapse-free survival in TNBC tumors.

## Summary points

- Flux activities could be used to characterize differences in metabolic pathways between groups of tumors and to associate them with clinical outcomes.
- There are differences in flux activities of nucleotide interconversion and ROS detoxification between breast cancer subtypes.
- Flux activities of vitamin A, beta-alanine metabolism and tetrahydrobiopterin metabolism predict distant metastasis-free survival in breast cancer patients.
- Flux activities of glycolysis and glutamate metabolism may predict distant relapse-free survival in TNBC.

## Ethical conduct

The study was approved by the Ethical Committee of Hospital 12 de Octubre and Hospital La Paz.

## Figure and table legends

Table 1: Patient characteristics.

Table 2: Weights assigned to each metabolic pathway contained in the predictor signature for all tumors.

Table 3: Weights assigned to each metabolic pathway contained in the predictor signature for TNBC tumors.

